## Supplementary figures and images for "Convergent evolution of aerobic fermentation through divergent mechanisms acting on key shared glycolytic genes"

### gal4_PTDH-UASG_GFP_DIC_R_p00_0_A01f42d3.TIF

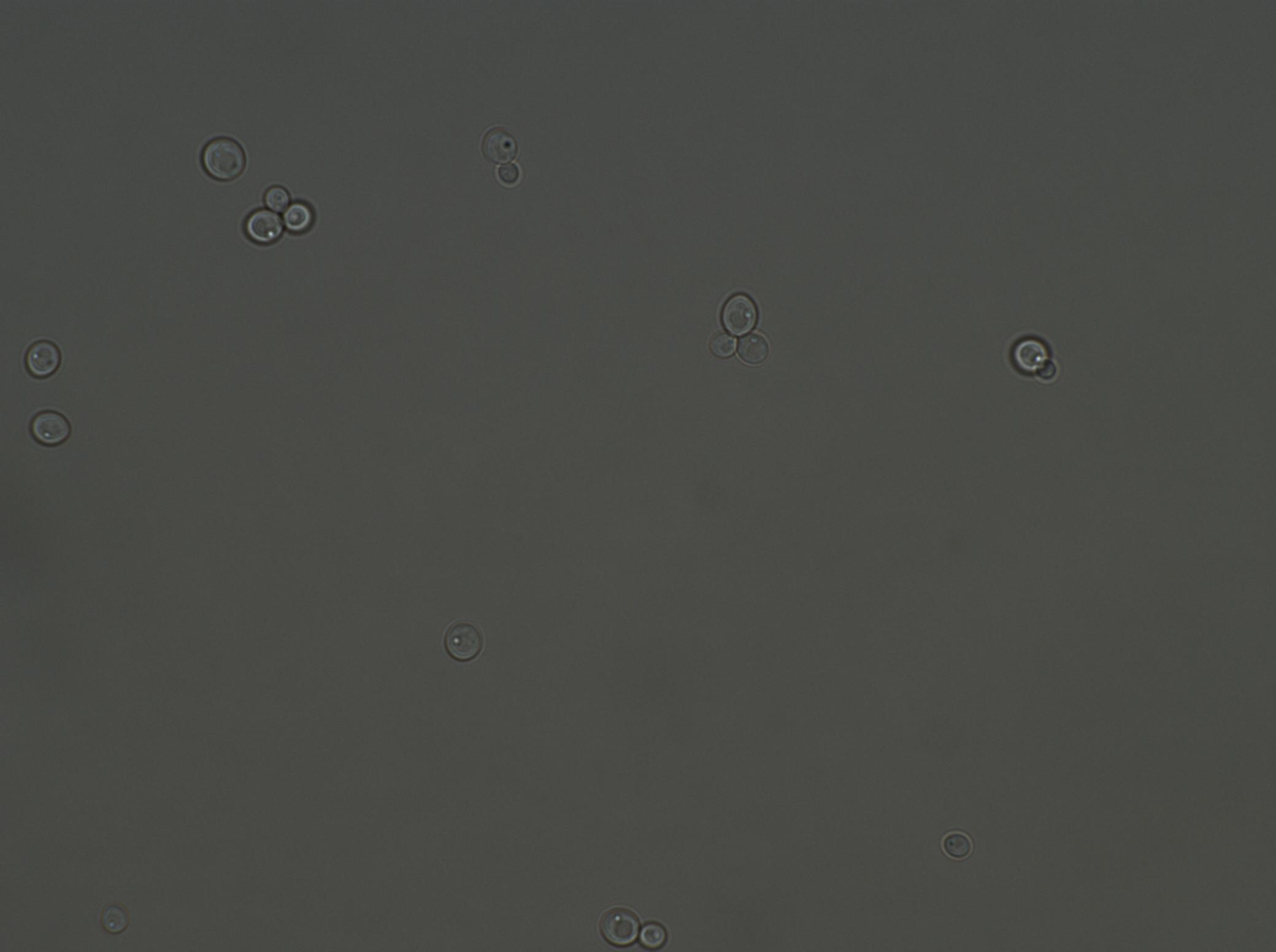

### gal4_PTDH-UASG_GFP_R_p00_0_A01f42d0.TIF

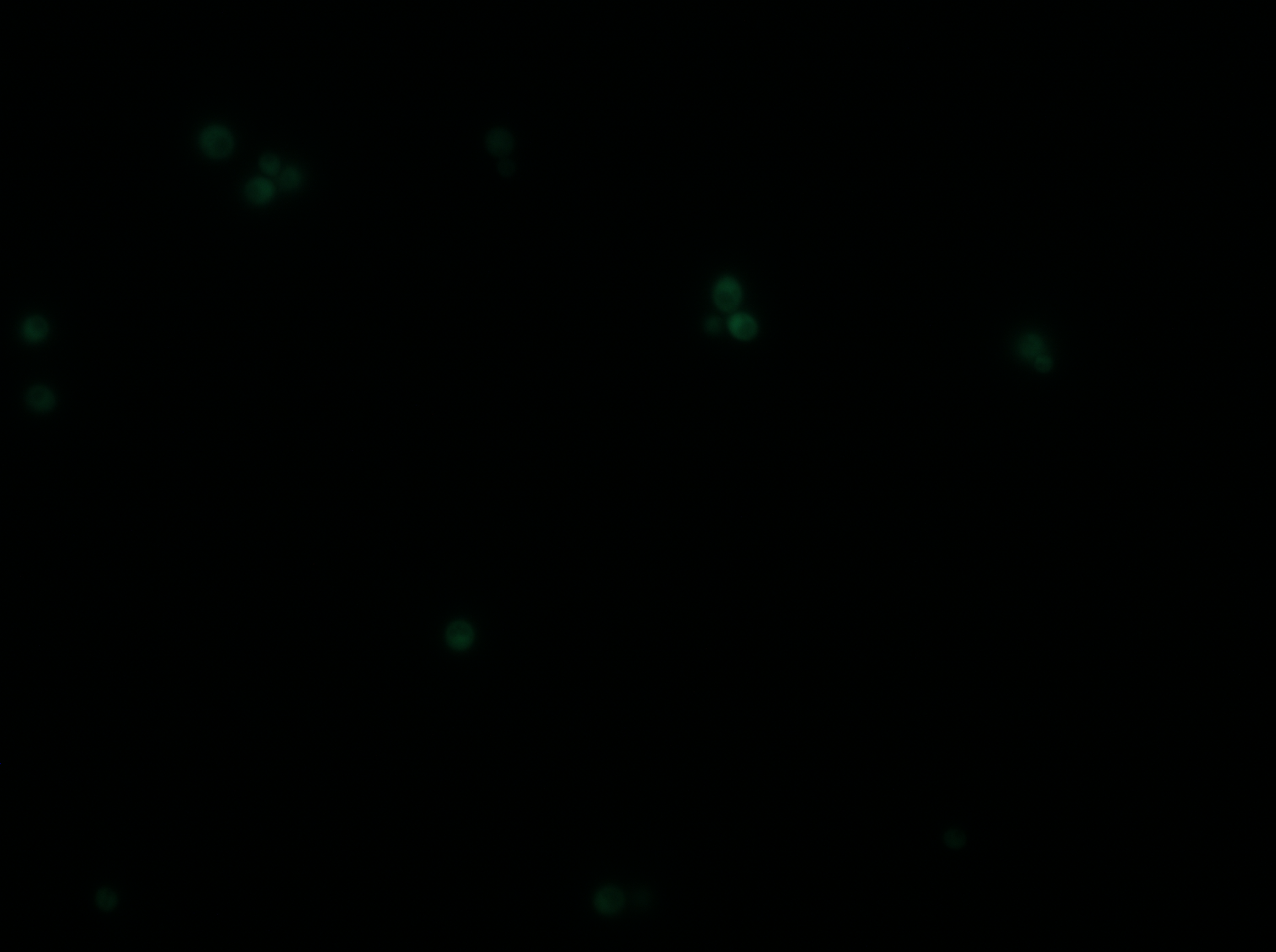

### gal4_PTDH_GFP_DIC_R_p00_0_A01f38d3.TIF

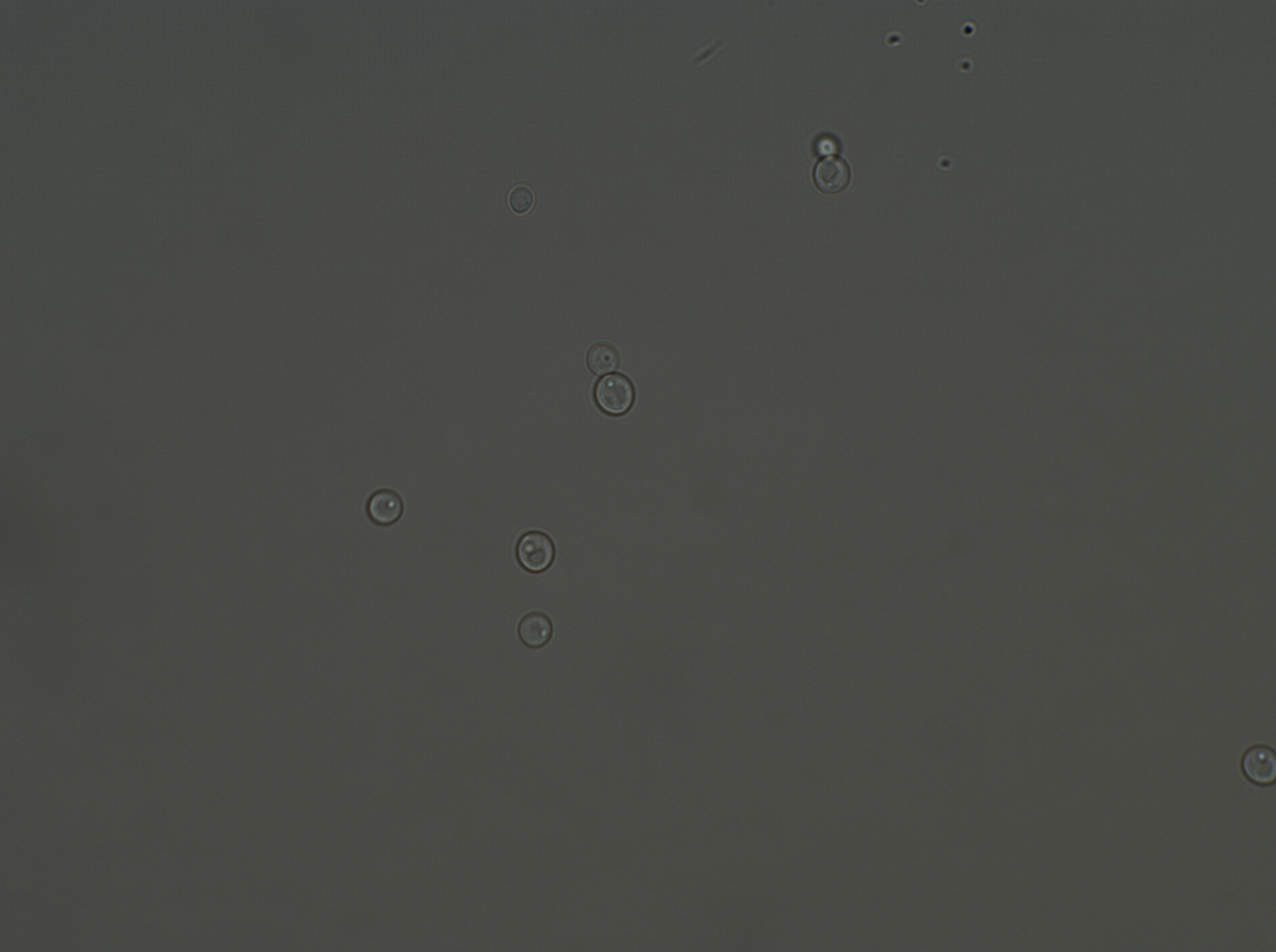

### gal4_PTDH_GFP_R_p00_0_A01f38d0.TIF

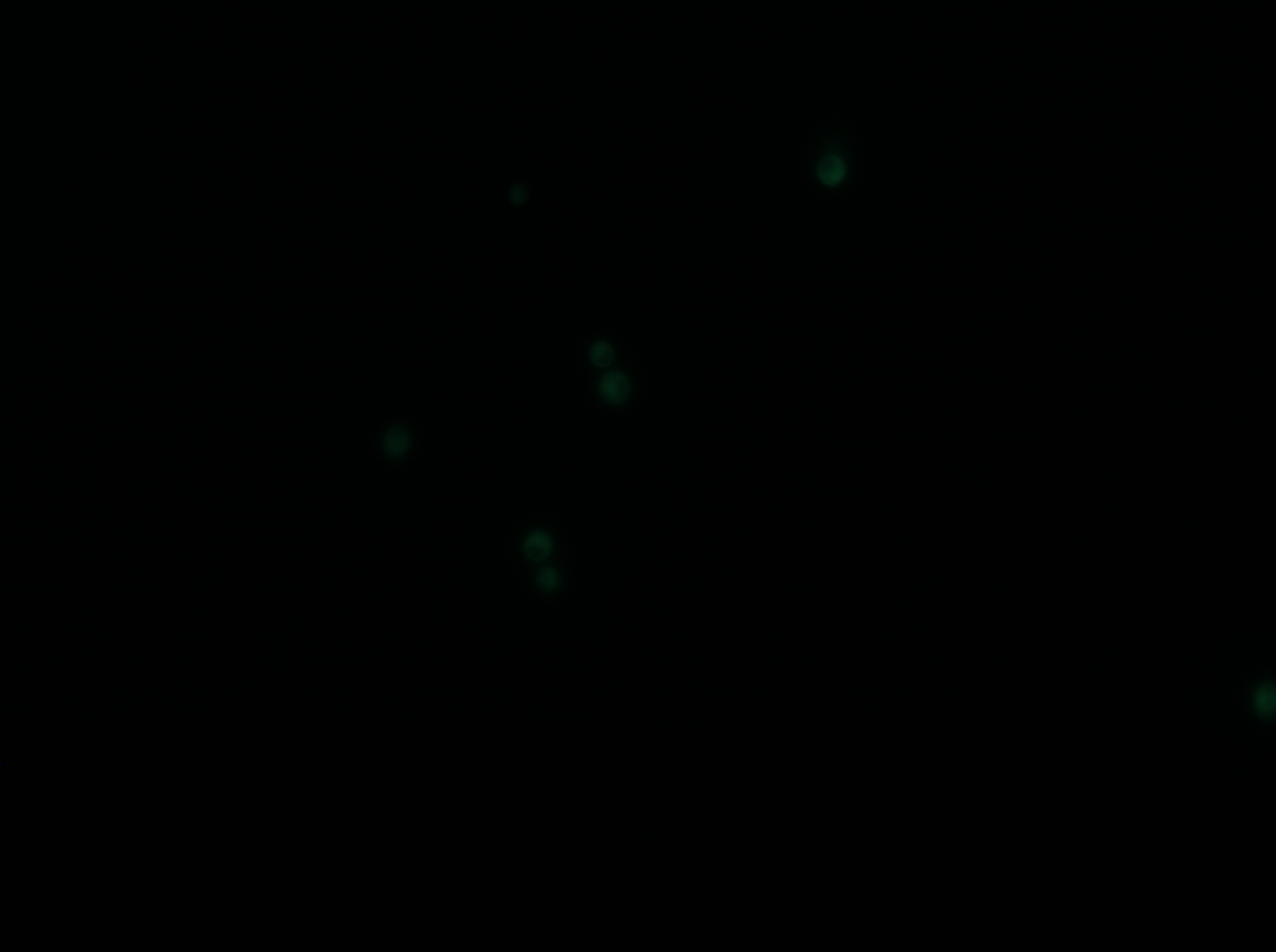

### WT_PTDH-UASG_GFP_DIC_R_p00_0_A01f17d3.TIF

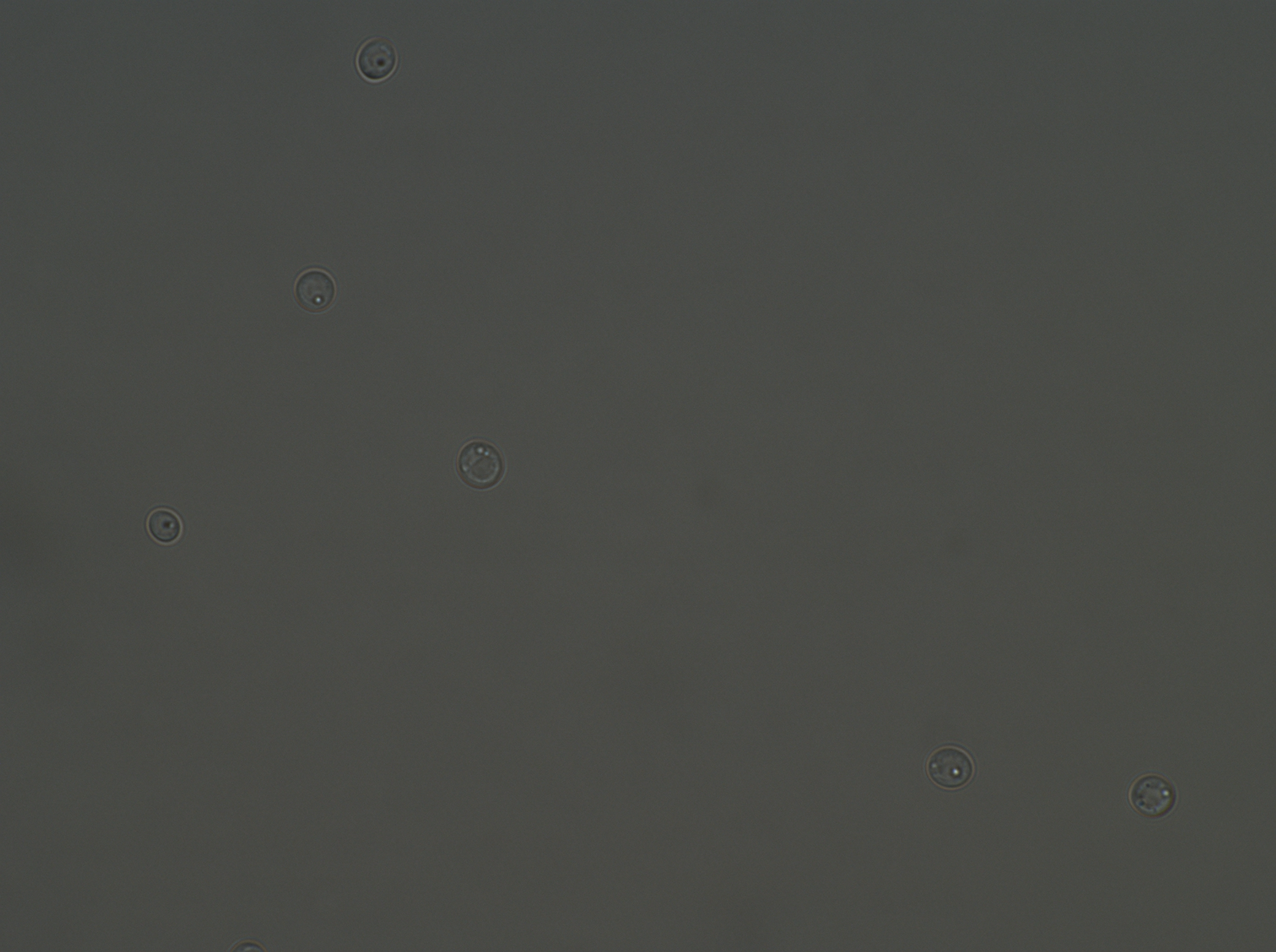

### WT_PTDH-UASG_GFP_R_p00_0_A01f17d0.TIF

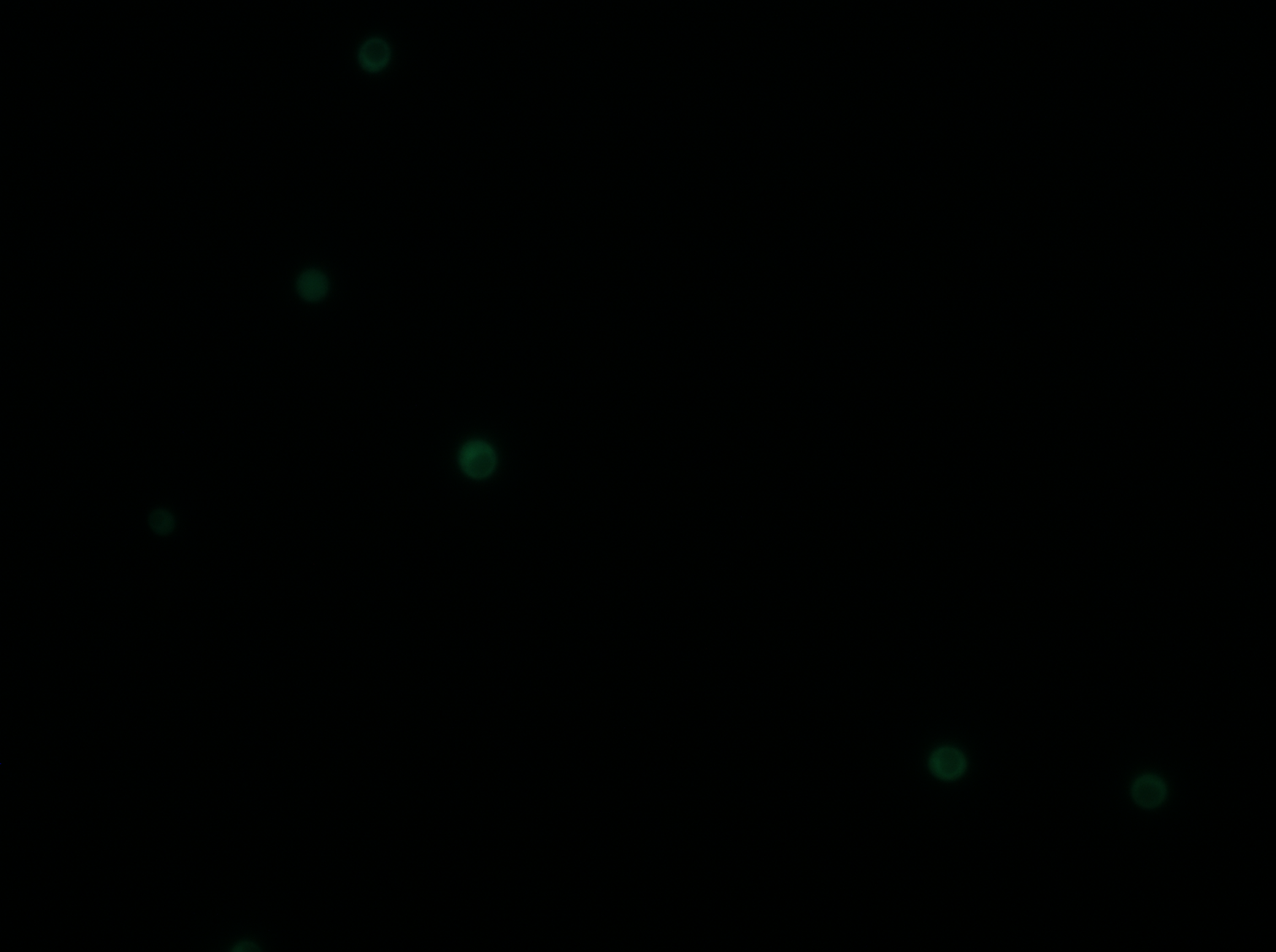

### WT_PTDH_GFP_DIC_R_p00_0_A01f03d3.TIF

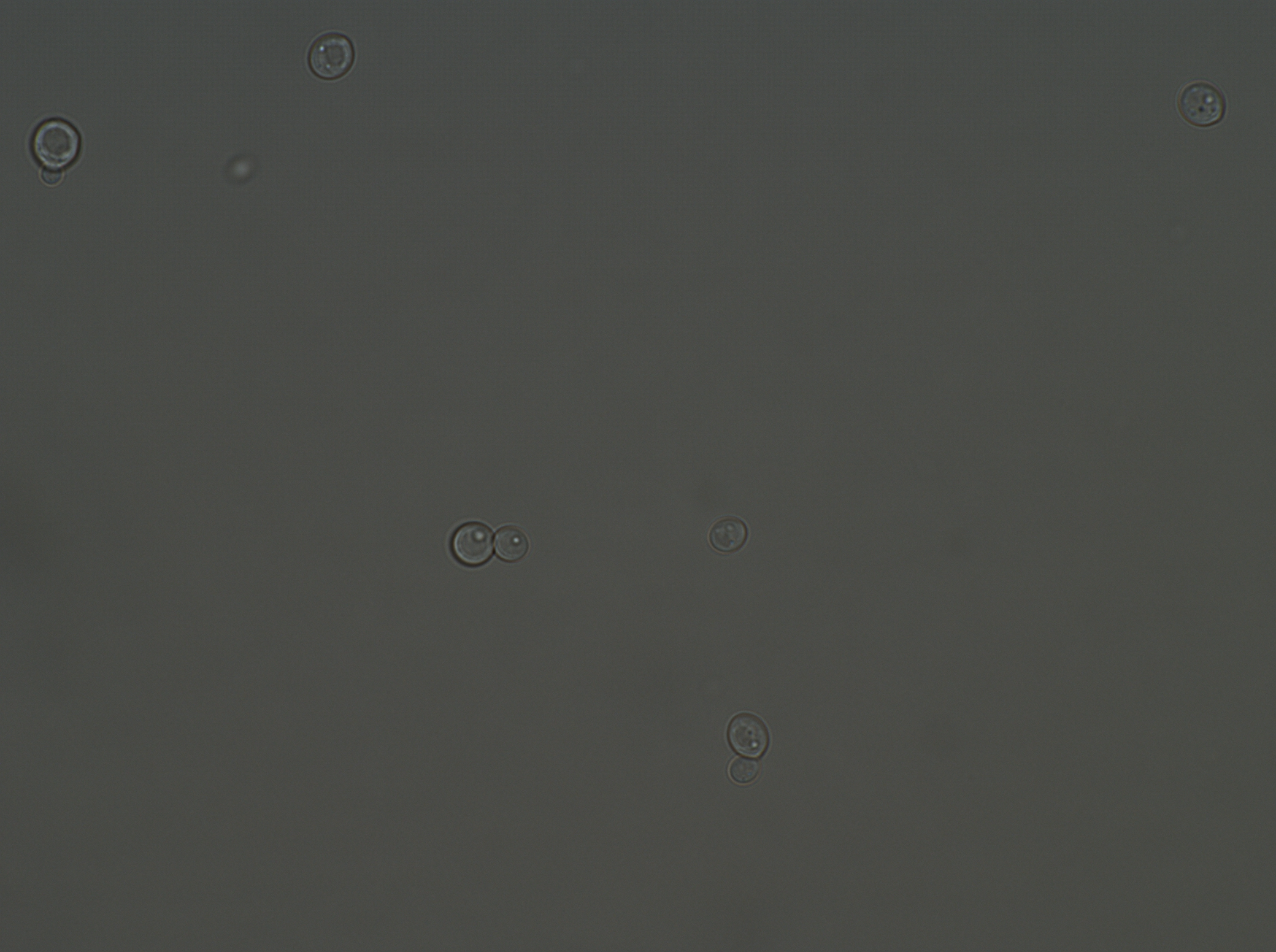

### WT_PTDH_GFP_R_p00_0_A01f03d0.TIF

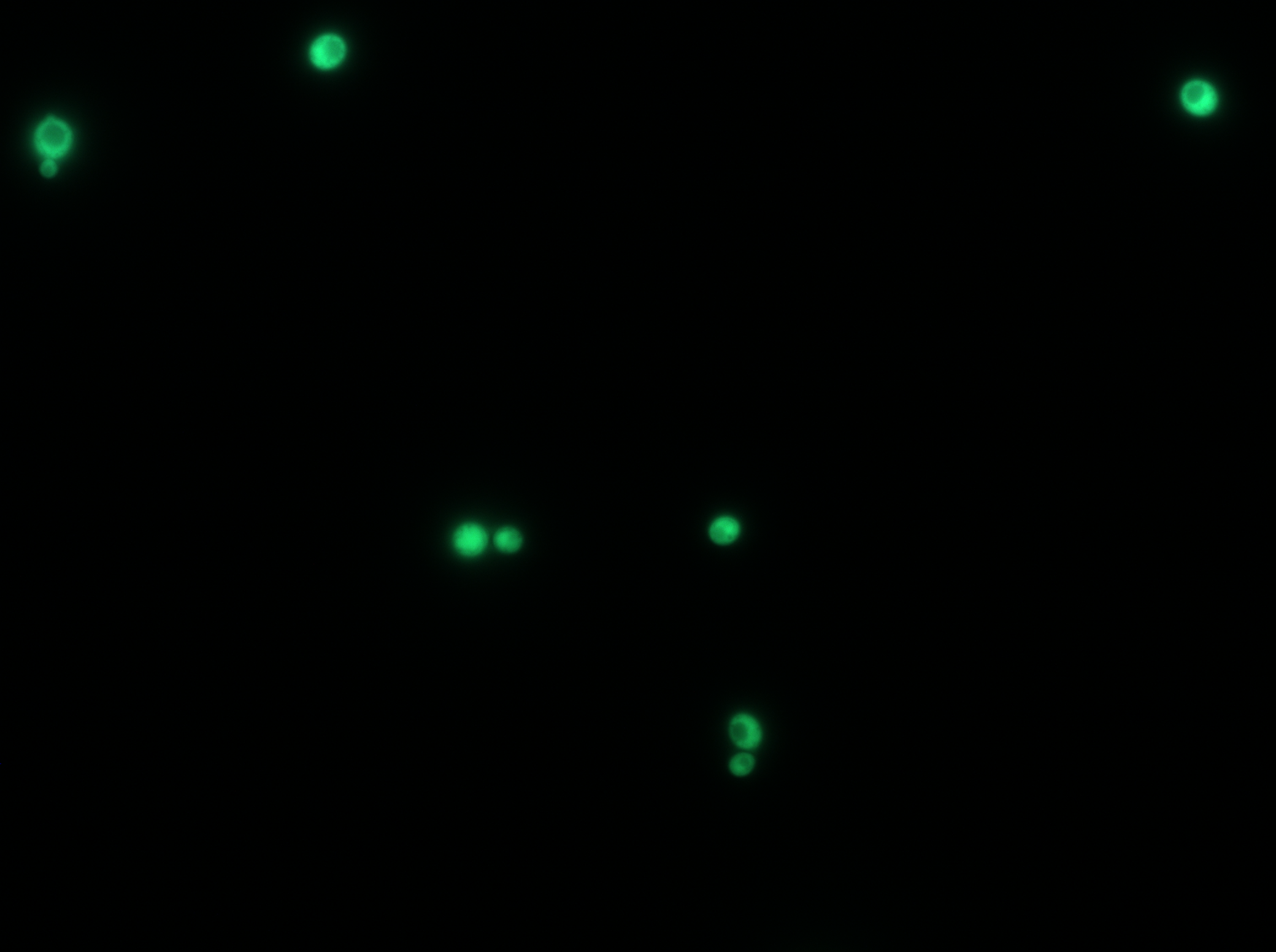
